# Reproducible data integration and visualization of biological networks in R

**DOI:** 10.1101/2022.04.15.488519

**Authors:** Florian Auer, Hryhorii Chereda, Júlia Perera-Bel, Frank Kramer

## Abstract

**Motivation:** Collaborative workflows in network biology not only require the documentation of the performed analysis steps but also of the network data on which the decisions were based. However, replication of the entire workflow or tracking of the intermediate networks used for a particular visualization remains an intricate task. Also, the amount and heterogeneity of the integrated data requires instruments to explore and thus comprehend the results.

**Results:** Here we demonstrate a collection of software tools and libraries for network data integration, exploration, and visualization to document the different stages of the workflow. The integrative steps are performed in R, and the entire process is accompanied by an interchangeable toolset for data exploration and network visualization.

**Availability:** The source code of the performed workflow is available as R markdown scripts at https://github.com/frankkramer-lab/reproducible-network-visualization. A compiled HTML version is also hosted on Github pages at https://frankkramer-lab.github.io/reproducible-network-visualization.

## 1 Introduction

Scientific research faces the problem of the reproducibility of the methods used in its various domains, with the effect that the reported results could not be reconstructed (Goodman et al., 2016). Using a steadily growing number of tools and working in multidisciplinary teams increasingly complicates replication, and necessitates to report not only the results but rather the entire workflow (Committee, 2021).

Analyses performed in network biology are no exception and require networks and their visualizations not to be seen only as input or result of the process. Networks are gradually enriched with additional data from various sources and the attention is mainly set on the applied methods. Intermediate networks are omitted although those could be utilized to document the progress and illuminate the contained information.

Furthermore, the visualizations of the integrated networks in any stage face the problem of choosing the appropriate attributes to focus on. Especially in a collaborative process this hampers the communication of relevant aspects in different steps of the workflow. Also, visual representations are subjective to the creator and may hide important features crucial for the understanding for collaborators and subsequent decisions for progression. An interactive exploration of the enriched networks can help to expose otherwise invisible characteristics but requires a seamless integration into the process.

Here we present a collection of tools to construct a reproducible workflow for network data integration and visualization, including the documentation of intermediate and final results. We demonstrate the workflow on previous results for the generation of patient-specific subnetworks for a large breast cancer data set (Chereda et al., 2021). Thereby we point out several options of software tools for the visualization that can be used individually or in combination to foster the reproducibility of the workflow.

## 2 Methods

### 2.1 Gene expression and molecular networks data

#### 2.1.1 Breast cancer data set

The studied breast cancer data set is composed from 10 microarray data sets publicly available at the Gene Expression Omnibus (GEO) (Barrett et al., 2013) repository by the accession numbers GSE25066, GSE20685, GSE19615, GSE17907, GSE16446, GSE17705, GSE2603, GSE11121, GSE7390, GSE6532. The expression data was measured on Affymetrix Human Genome HG-U133 Plus 2.0 and HG-U133A arrays and previously preprocessed and studied (Bayerlová et al., 2017), including the prediction of the molecular subtypes for the Breast cancer samples. Sample selection, filtering, combination and normalization was performed according to previous work (Chereda et al., 2021) and resulted in a data set containing 12,179 genes in 969 patients. The patients’ metastatic status were derived from the occurrence of distant metastasis within the first 5 years (393 patients) or absence with the last follow-up between 5 and 10 years (576 patients).

#### 2.1.2 Protein interaction networks

The protein-protein interaction (PPI) network from the Human Protein Reference Database (HPRD) (Keshava Prasad et al., 2009; Mishra et al., 2006; Peri et al., 2003) was used as basis for capturing the relations between the expressed genes. The information contained in molecular network is based on evidence from in vitro and in vivo yeast two-hybrid analyses and constitutes of undirected binary interactions between pairs of proteins.

The disconnected graph consists of 9,898 vertices which decreased after mapping the genes of the breast cancer data set onto the PPI network to 7,168 vertices in 207 connected components. For further analysis only the main connected component was used, which consisted of 6,888 vertices, while the remaining components only contained 1 to 4 vertices. The main reason for this choice was that the Graph-CNN algorithm requires a connected graph as input.

### 2.2 Data processing

#### 2.2.1 Relevance score

The computation of a patient-specific relevance score is a two-step process: Firstly, a Graph Convolutional Neural Network (Graph-CNN) is trained on the gene expression and molecular network data to predict the metastatic status for a patient. Secondly, the Graph Layer-wise Relevance propagation (GLRP) algorithm is applied to a patient’s prediction to determine the relevance of the genes to the predictive outcome. (Chereda et al., 2021)

The Graph-CNNs were trained on the gene expression dataset with the HPRD PPI network as prior knowledge with a 10-fold cross validation over a whole dataset to estimate the predictive performance of Graph-CNN. For the generation of the relevance scores, the gene expression dataset was randomly split in training (90%) and test (10%) set. The Graph-CNN was trained using manually selected hyperparameters from 10-fold cross validation, and subsequently used to predict metastatic events for the test set consisting of 97 patients.

For the subsequent analysis, meaning the generation of patientspecific subnetworks, only 79 patients were considered with matching predicted and reported metastasis. The GLRP method was applied for those patients to determine the relevance of the genes to the prediction, thus called relevance score.

#### 2.2.2 Molecular tumor board report analysis

Actionable genes present in the patient-specific subnetworks were identified using the Molecular Tumor Board (MTB) report (referred to as “MTB report”) methodology described in Perera-Bel et al. Therefore, the algorithm was extended by inferring gain of function alterations from high expression, and loss of function alterations from low expression respectively. The gene expression levels were derived based on the gene expression throughout the whole patient cohort with the 25% and 75% quantiles as boundaries for low, normal and high levels of expression.

Although information about specific gene variants is not present in the breast cancer gene expression data set due to the used quantification method, the results can be used to define specific panels for subsequent sequencing.

### 2.3 Data integration and modeling

#### 2.3.1 The NDEx platform

The distribution of biological networks is an important aspect in collaborative workflows. Working on the same data basis can be challenging to be arranged, be it providing the networks used as resource for the initial analysis or sharing intermediate and final results.

The Network Data Exchange (NDEx) (Pillich et al., 2017; Pratt et al., 2015) is an online commons specifically designed for the exchange of and collaboration on biological networks. Networks can be uploaded and shared with individual persons or groups and remain visible only to this peer to prevent premature disclosure of this valuable information.

For the publication of the results, NDEx then can be used to propagate the network data supplementary to manuscripts, and as possible resources for further analyses. Furthermore, a comprehensible collection of networks is publicly available on the platform as for example the NCI Pathway Interaction Database (PID) (Schaefer et al., 2009) from which NDEx initially originated.

#### 2.3.2 ndexr

Programmatic interaction with the NDEx platform from within R (R Development Core Team, 2008) is provided by the *ndexr* package (Auer et al., 2018). The package enables the search of public and private networks on the platform as well as the exchange of networks with the platform. Moreover, it provides functions to adjust the accessibility and visibility of the networks as well as options for sharing with specific users and groups.

In this work we use *ndexr* to document the progress of the integrative network analysis. The HPRD PPI network available at the NDEx platform was retrieved, and the intermediate stages from integration of the gene expression with the network to the creation of the single patientspecific subnetworks are saved using this package.

#### 2.3.3 RCX

The NDEx platform uses for the exchange of the network data their proprietary Cytoscape Exchange (CX) data format. It is an aspect oriented and JSON based data structure tailored to the transmission of biological networks. It utilized established web standards for the transmission and thereby encapsulates the different components of the network (i.e., nodes, edges, layout and visual representation, and associated attributes) into separated modules (aspects). The different aspects are independent by itself but can refer to each other, if necessary, for example edges refer to the nodes they connect, and the cartesian layout to the nodes to which they assign the position.

CX originated from the cooperation with the Cytoscape consortium and consequently inherited aspects dedicated to capture the visual representation of the network. Moreover, one remarkable feature of CX in contrast to other network formats is that the visual representation is a part of the network itself.

The *RCX* package (Auer & Kramer, 2022) includes functions and models to facilitate working with biological networks in CX format within R. The RCX data model, including separate models for the single aspects, is thereby the adaptation of CX to standard R data types and structures. Due to the fundamental differences between the table-based view on data in R and the object-oriented composition in JSON, and hence CX, the *RCX* package offers specialized functions for conversion and handling of the networks. Besides the lossless conversion to the CX format also *igraph* (Csardi & Nepusz, 2006) and Bioconductor *graph* (Gentleman et al., 2021) are supported, both established libraries for graph analysis and visualization. Furthermore, *RCX* includes functions for the creation, modification and validation of networks in this format to facilitate usability.

#### 2.3.4 Generation of patient-specific subnetworks

The *ndexr* package uses *RCX* as data model for the integration of the gene expression values and levels, and relevance scores with the HPRD PPI network to generate the subnetworks, as well as to capture and store the visualizations (Fig. 1). The resulting integrated network is available on the NDEx (UUID 833b1cee-42f6-11ec-b3be-0ac135e8bacf) and forms the basis for the subnetwork generation.

**Fig. 1.**
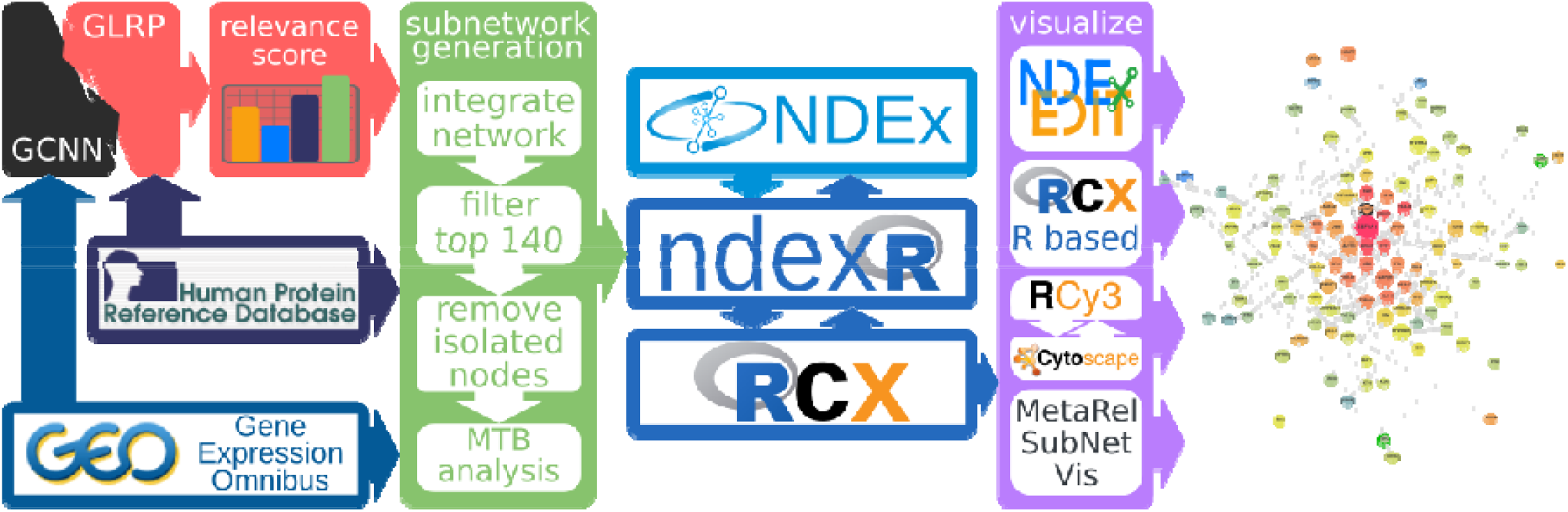
Network data integration and visualization workflow. HPRD and GEO serve as raw data resource for the generation of patient-specific subnetworks and used for calculation of the relevance scores. The networks are stored on the NDEx platform using *ndexr* and handled by *RCX* which was also used for the visualization, additionally to *NDExEdit, MetaRelSubNetVis,* and RCy3 and Cytoscape.

For each of the 79 patients the 140 most relevant genes were used to induce the intermediate patient-specific subnetworks. Those subnetworks then were combined and again only the main connected component consisting of 407 nodes used. For these final patient-specific subnetworks the MTB report analysis was applied and the results integrated into the combined network (UUID a420aaee-4be9-11ec-b3be-0ac135e8bacf), which then was used as basis for the different visualization approaches. The patient-specific subnetworks are also individually accessible at the NDEx platform within a network collection (UUID 5d308fbb-42da-11ec-b3be-0ac135e8bacf).

Since the single integration steps build upon each other it is necessary to track the preceding networks for reproducibility of the integration steps. Therefore, the source networks are listed by their UUID within the network attributes of the current network model.

## 3 Results

### 3.1 MetaRelSubNetVis

*MetaRelSubNetVis* is a web-based tool for the interactive exploration of integrated patient subnetworks and comparison of those networks between patient groups. The combined integrated subnetwork is directly loaded from NDEx and visualized within the web-browser with a concentric layout applied by default. The visual representation of the subnetworks can be adjusted to base on the integrated data. The included genes can be sized and colored using a gradient for expression or relevance score values. Alternatively, different colors for the expression levels can be set. Additionally, the results of the MTB analysis can be highlighted within the graph as well as selected nodes.

The different patient groups, i.e., metastatic and non-metastatic, can be visualized and compared within the same graph. For a selected patient of each group the nodes are split and colored by the value of the patient of the corresponding group (Fig. 3). Shared genes are sized greater than individual ones and to put even more emphasis on common nodes only those can be shown.

The visualization can be explored interactively by moving nodes or adjusting the thresholds for relevance scores or gene expression to display the nodes. Thereby, the position of the nodes is preserved between visualization and patient selection changes to facilitate identification of the same entities across the different subnetworks.

Furthermore, custom links can be generated that lead directly to the selected patients and their visualization. These links can be used to communicate and reference specific findings within the subnetworks. They can also be provided along publication for illustration and be embedded on websites for reference or interactive exploration (Fig. 2).

**Fig. 2.**
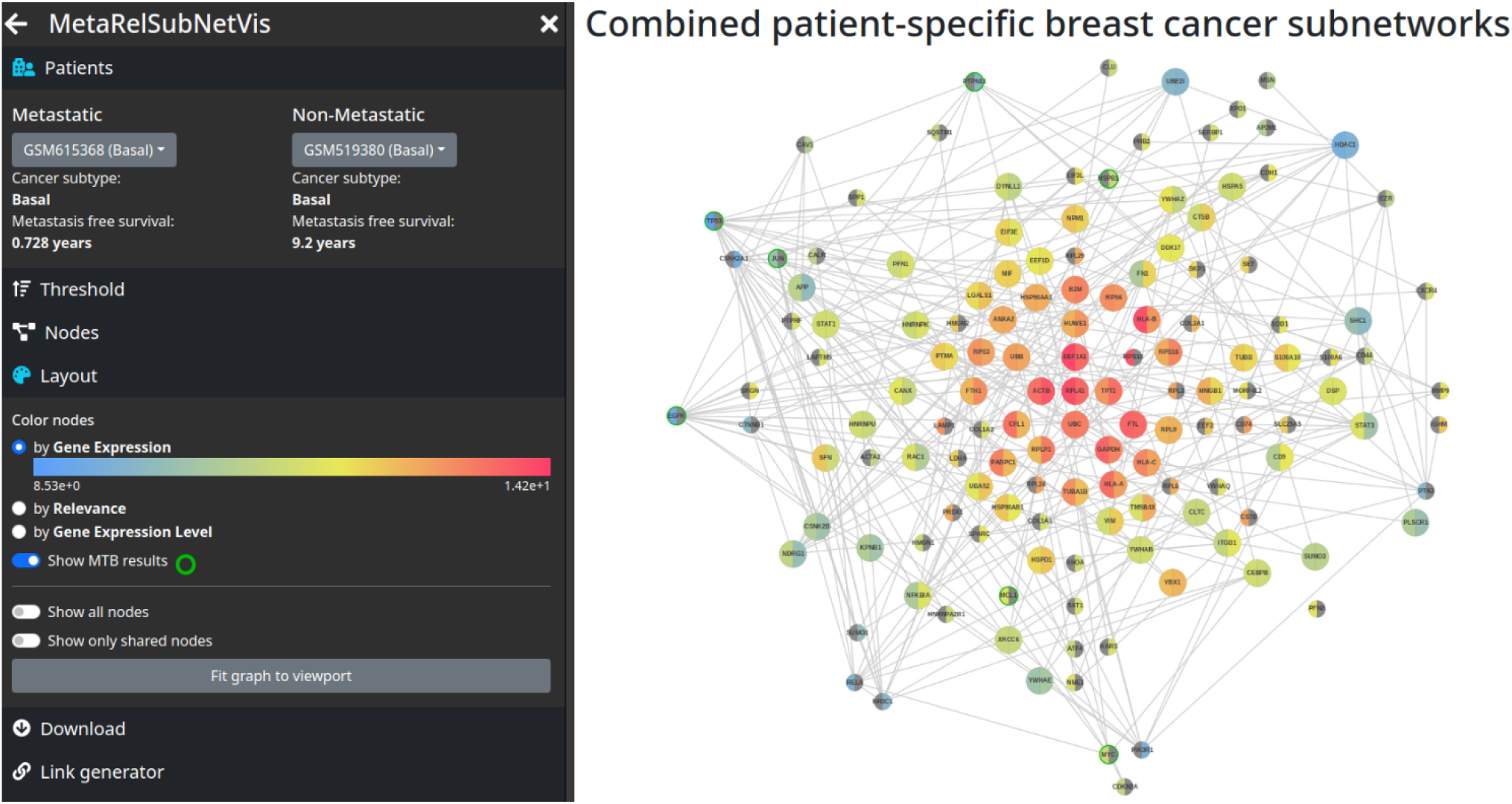
Comparative visualization of patient GSM615368 and GSM615184 on MetaRelSubNetVis colored by gene expression. The visualization network is available at https://frankkramer-lab.github.io/MetaRelSubNetVis?uuid=a420aaee-4be9-11ec-b3be-0ac135e8bacf&pa=GSM615368&pb=GSM519380&th_GE=8.53234588826455&th_Score=0.00029828155&col=GE&size=GE&all=false&shared=false&bool=MTB&sb=0&cP=0&cT=1&cN=1&cL=0&cD=1&cG=1&cIm=1&bb=true

### 3.2 NDExEdit

The visualization of networks, even only simple ones, often requires additional software to be installed on the local machine. *NDExEdit* is a web-based approach for the data-dependent visualization of networks where those can be loaded directly from the NDEx platform. For a loaded network the single attributes, and their distribution, can be explored and visual attributes applied based on the contained data. The network can be arranged, different layouts applied, and the results saved within the network. This can be used to highlight for example the number of occurrences of the different genes across the combined network (Fig. 4), or to define the same visual styles for a single patient as demonstrated with *MetaRelSubNetVis.* Finally, the resulting networks with included visualization can directly exported to the NDEx platform, downloaded as CX file, or exported as publication-ready images in various formats.

**Fig. 3.**
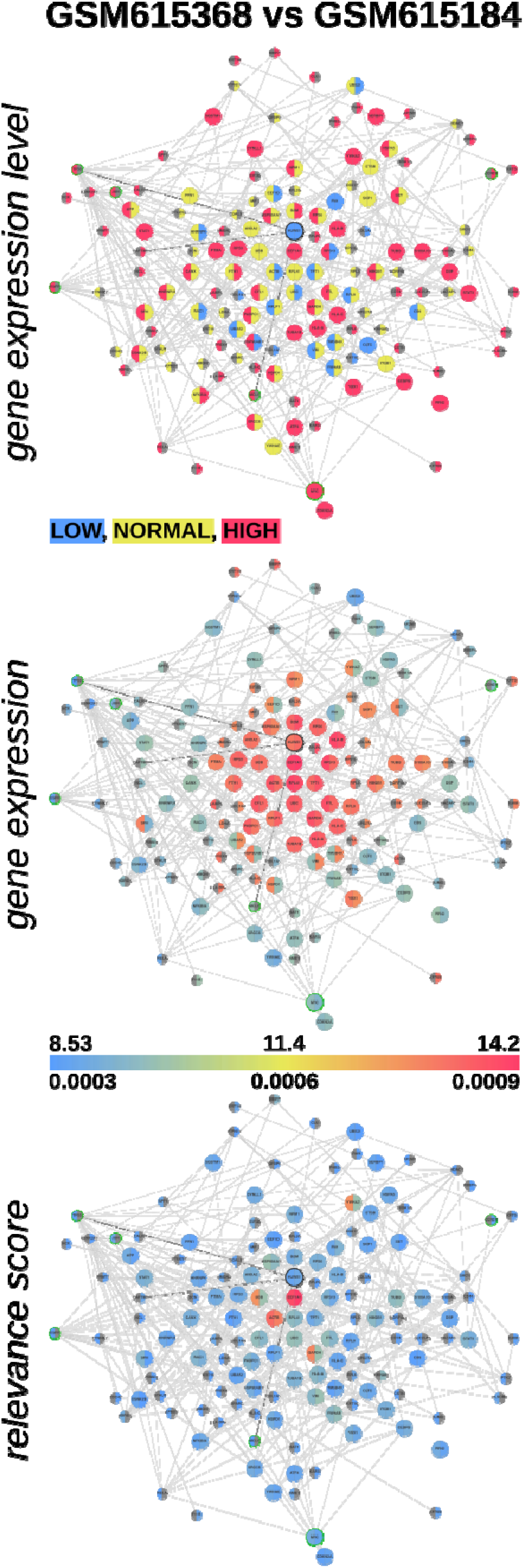
Comparative visualization of patient GSM615368 and GSM615184 colored by gene expression level and value, and relevance score.

**Fig. 4.**
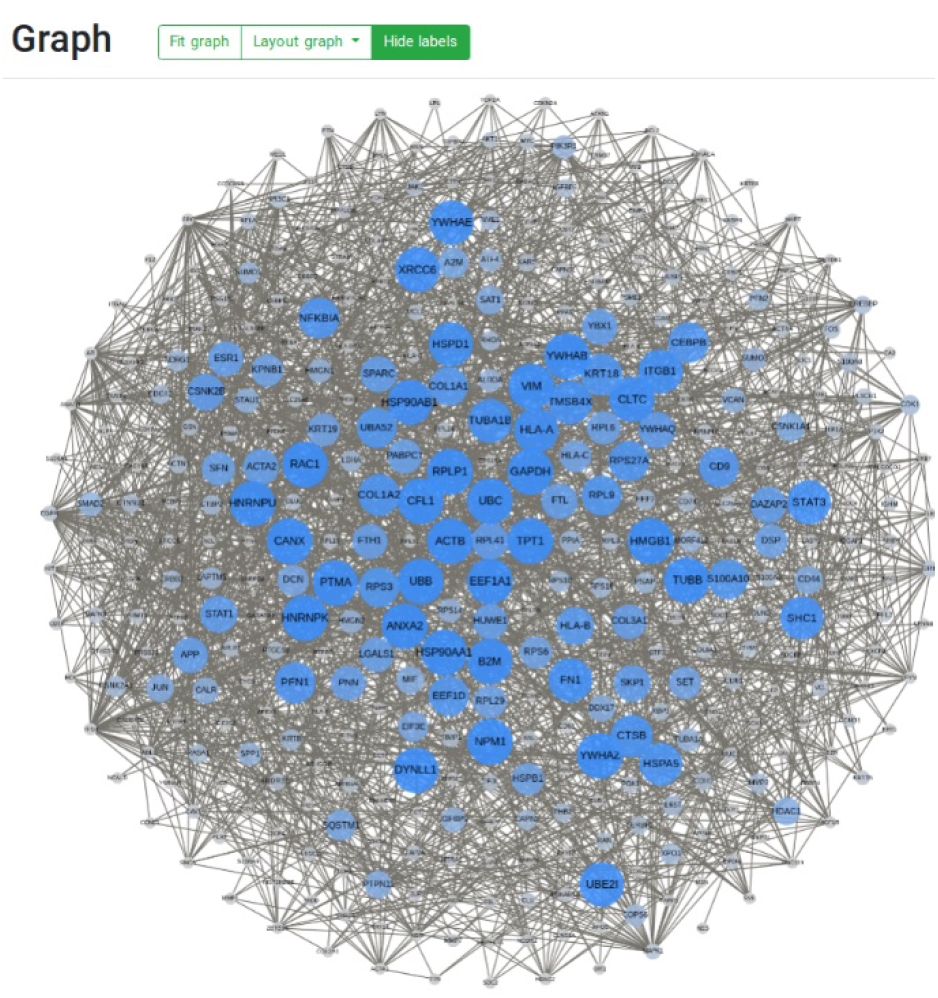
Combined patient-specific subnetworks. The nodes are colored and sized by the number of occurence across the 79 patients. The network is available on the NDEx platform by the UUID a420aaee-4be9-11ec-b3be-0ac135e8bacf.

### 3.3 R based visualization

#### 3.3.1 igraph

Within the R environment the most prominent software package for graph manipulation, analysis, and visualization is the *igraph* library. The package thereby follows the typical R methodology of defining the visual attributes for each node and edge individually and explicitly before plotting the graph. This way similar visualizations can be created as with the above tools, but the attribute values for single nodes (and edges) must be set individually. Also, the sharing and exploration of networks in this format is rather inconvenient and especially the latter requires expertise with appropriate tools.

#### 3.3.2 Cytoscape and RCy3

One of the most widely used software tools for the visualization of biological networks is Cytoscape (Shannon et al., 2003). It allows the import of networks in various formats, including CX from file or directly from the NDEx platform. Besides the simple definition of visual properties Cytoscape offers plenty of tools for network analysis, that can be extended even further by custom plugins.

In contrast to the *igraph* package where individual values are assigned to the nodes and edges, Cytoscape defines mappings based on attributes of those. This not only allows a more generalized definition of the visual properties, but also promotes the reuse of the created visual styles for different networks.

Cytoscape provides a REST API which can be used with the R package *RCy3* (Gustavsen et al., 2019) to access the software in a programmatic manner. This allows to remotely control Cytoscape to reproduce the visualization of the patient-specific subnetworks with the same representation as in NDExEdit. Since both tools allow the export of the visualized networks to the NDEx platform the visualizations can be continued or refined in both tools interchangeably.

#### 3.3.3 RCX

The *RCX* package was not only used for handling of the network data and integration steps, but also to define and apply layouts and visual attributes. Therefore, the package includes functions to produce visualizations of the networks consistent with those on the NDEx platform, on NDExEdit and within Cytoscape. This consistency is based on the aspect-oriented structure of the RCX data model which includes the properties for the visual representation. We show how the aspect for the visual representation can be created from scratch, including the necessary properties, mappings and dependencies for the nodes and edges.

However, for users unfamiliar with the Cytoscape visual properties this approach is arduous. Therefore, we demonstrate a simpler strategy by reusing the visualization created with *NDExEdit* for a single patient. The visual properties of the downloaded network are adjusted for the remaining patients for mapping the corresponding patient data. Subsequently the patient-specific subnetworks including their visualizations are exported the NDEx platform.

## 4 Discussion

When working on data integration with biological networks in R the most straight forward approach is to use the most established *igraph* library, especially if it requires methods for graph and network analysis. However, visualization and distribution of the integrated network models is rather limited. Tools like Cytoscape simplify the visualization process of the created networks, but still require a solution for network import and distribution.

The NDEx platform offers a solution for management and collaboration and is already integrated into Cytoscape. Together with the *RCy3* package they provide an option to load, visualize and store and share the networks. Nevertheless, this may constitute an unnecessary detour, especially if the visualization is performed by another party. The usage of the *RCX* package for network integration, or through conversion from the *igraph* models provides in combination with *ndexr* a shortcut to the NDEx platform.

The RCX package itself can be used for the visualization of networks but instead of manually defining the visual properties its greater benefit lies in the reuse and adjustment of those. Again, Cytoscape can be used for the creation of visual properties but NDExEdit on the contrary does not require installation of the software. Furthermore, it provides options to explore the contained data and adjust the visual mappings accordingly which supports especially users unfamiliar with the network. This though comes with the cost of NDExEDIT relying on the NDEx platform due to missing support of import options for other network formats.

Although the NDEx platform promotes sharing and referencing the resulting networks it still has its shortcomings in terms of the representation of integrated network data. *MetaRelSubNetVis* offers an addition to NDEx by allowing to interactively explore the networks with a consistent network structure, a property-based visualization, and a groupwise comparison. Furthermore, the specific visualization settings can be shared additionally to the network. However, this requires the information about the visualization parameters to be included within the network.

## 5 Conclusion

Here we presented different approaches for the reproducible network data integration and visualization. The presented tools constitute of established software and libraries each with its own advantages and usecases. They mainly evolve around the NDEx platform which enables storage and distribution of results of an analysis and allows the documentation of the performed steps. Together with the inclusion of the visualization within the networks it not only contributes to the comprehensibility of the results but also fosters their reproducibility.

The application for the generation of patient-specific subnetworks illustrates its applicability in a typical bioinformatics workflow. The proposed solutions are not exclusive but rather complementary to established methods and demonstrate their benefits especially through flexibility in their usage. The visualization of intermediated network results brings additional insights to the performed integration steps. Only the combination of the here discussed software tools, platforms and packages promotes an environment for the reproducibility network data integration and accompanying visualization.

